# *In vivo* structural characterization of the whole SARS-CoV-2 RNA genome identifies host cell target proteins vulnerable to re-purposed drugs

**DOI:** 10.1101/2020.07.07.192732

**Authors:** Lei Sun, Pan Li, Xiaohui Ju, Jian Rao, Wenze Huang, Shaojun Zhang, Tuanlin Xiong, Kui Xu, Xiaolin Zhou, Lili Ren, Qiang Ding, Jianwei Wang, Qiangfeng Cliff Zhang

**Affiliations:** MOE Key Laboratory of Bioinformatics, Beijing Advanced Innovation Center for Structural Biology & Frontier Research Center for Biological Structure, Center for Synthetic and Systems Biology, Tsinghua-Peking Joint Center for Life Sciences, School of Life Sciences, Tsinghua University, Beijing, 100084, China; Center for Infectious Disease Research, School of Medicine, Tsinghua University, Beijing, 100084, China; NHC Key Laboratory of Systems Biology of Pathogens and Christophe Mérieux Laboratory, Institute of Pathogen Biology, Chinese Academy of Medical Sciences & Peking Union Medical College, Beijing, 100730, China; Key Laboratory of Respiratory Disease Pathogenomics, Chinese Academy of Medical Sciences and Peking Union Medical College, Beijing, 100730, China

## Abstract

SARS-CoV-2 is an RNA virus of the *Coronaviridae* family that is the causal pathogen of the ongoing Coronavirus Disease 2019 pandemic. There are currently no antiviral drugs or vaccines to treat COVID-19, and the failure to identify effective interventions can be blamed on our incomplete understanding of the nature of this virus and its host cell infection process. Here, we experimentally determined structural maps of the SARS-CoV-2 RNA genome in infected human cells and also characterized *in vitro* refolded RNA structures for SARS-CoV-2 and 6 other coronaviruses. Our *in vivo* data confirms several structural elements predicted from theoretical analysis and goes much further in revealing many previously unknown structural features that functionally impact viral translation and discontinuous transcription in cells. Importantly, we harnessed our *in vivo* structure data alongside a deep-learning tool and accurately predicted several dozen functionally related host cell proteins that bind to the SARS-CoV-2 RNA genome, none of which were known previously. Thus, our *in vivo* structural study lays a foundation for coronavirus RNA biology and indicates promising directions for the rapid development of therapeutics to treat COVID-19.

**HIGHLIGHTS:**

- We mapped the *in vivo* structure and built secondary structural models of the SARS-CoV-2 RNA genome
- We discovered functionally impactful structural features in the RNA genomes of multiple coronaviruses
- We predicted and validated host cell proteins that bind to the SARS-CoV-2 RNA genome based on our *in vivo RNA* structural data using a deep-learning tool

## INTRODUCTION

Coronavirus Disease 2019 (COVID-19), caused by a coronavirus named Severe Acute Respiratory Syndrome coronavirus 2 (SARS-CoV-2), has spread globally and brought devastating to public health and economic damage, with more than 10 million people infected and 500 thousand deaths (Dong E et al., 2020; Wu et al., 2020). Although huge global efforts have been devoted to understanding and fighting SARS-CoV-2—including extensive molecular virology studies examining the overall viral particle and viral protein structures (Gao et al., 2020; Lan et al., 2020; Walls et al., 2020; Yan et al., 2020), transcriptome architectures (Kim et al., 2020), and host cell-viral interactomes (Gordon et al., 2020), as well as mechanistic studies of the virus infection process and antiviral immune responses (Hoffmann et al., 2020; Ni et al., 2020; Walls et al., 2020)—there are presently no effective antiviral treatments or approved vaccines.

SARS-CoV-2 is an RNA virus of the *Coronaviridae* family, which also includes the SARS-CoV virus that caused the SARS outbreak in 2003 (Peiris et al., 2003) and the Middle East respiratory syndrome coronavirus (MERS-CoV) that caused the MERS outbreak in 2012 (Zaki et al., 2012). The genome of SARS-CoV-2 is an approximately 30kb, single-stranded, positive-sense RNA including a 5’ cap structure and a 3’ poly(A) tail. After cell entry, the viral genome can be translated into proteins directly, or can serve as the template for replication and transcription. During translation, SARS-CoV-2 produce nonstructural proteins (nsps) from two open reading frames (ORF1a and ORF1b), and a number of structural proteins from subgenomic viral RNAs. Upon production of negative-sense RNA by the nsp12 protein (an RNA-dependent RNA polymerase), both positive-sense genomic RNA and sub-genomic RNAs can be generated. The RNA comprising the SARS-CoV-2 genome is packaged by structural proteins encoded by sub-genomic RNAs.

It is notable that a large majority of molecular virology studies of SARS-CoV-2 (and indeed studies of most other viruses) have focused on viral proteins. For example, structural determination of the receptor-binding domain (RBD) of the spike protein of SARS-CoV-2 bound to the cell receptor ACE2 provided atomic details on the initial step of infection (Lan et al., 2020; Walls et al., 2020; Yan et al., 2020). Knowledge of SARS-CoV-2-human protein-protein interactions revealed the basis of how SARS-CoV-2 reshapes cellular pathways and identified druggable host factors for FDA-approved drugs and small compounds (Gordon et al., 2020). And tracking and analysis of changes in the key proteins of SARS-CoV-2 discovered an important mutation that is associated with increased transmission (Korber et al., 2020). These studies have been valuable for revealing mechanistic insights to deepen understanding of molecular virology and epidemiology and to aid development of antiviral therapeutics.

However, SARS-CoV-2 is an RNA virus; its RNA genome itself is a central regulatory hub for controlling and enabling its function. Recent advances in technologies and basic studies have emphasized that RNA molecules in cells fold into complex, higher-order structures that are integral to their cellular functions (Brion and Westhof, 1997; Piao et al., 2017). For viruses specifically, there are many examples of functional impacts from such RNA structural elements: studies of flaviviruses have confirmed that intramolecular RNA-RNA interactions between the 5’- and the 3’-UTRs promote genome circularization and help coordinate replication (de Borba et al., 2015; Nicholson and White, 2014); the structure of the internal ribosome entry site in 5’UTR is crucial for hepatitis C virus (HCV) translation (Fraser and Doudna, 2007; Kieft, 2008); and the multi-pseudoknot structures in the 3’ UTR of ZIKV and other flaviviruses have been shown to stall the RNA exonuclease Xrn1, thereby giving rise to sub-genomic flavivirus RNAs (sfRNAs) that help the virus evade cellular antiviral processes (Akiyama et al., 2016; Filomatori et al., 2017). However, despite functional characterization of several RNA structural elements of SARS (Robertson et al., 2005), and theoretical predictions recently available along with the sequences of SARS-CoV-2 (Andrews et al., 2020; Rangan et al., 2020), it is clear that a more comprehensive analysis of the structure of the SARS-CoV-2 RNA genome as it exists in a physiologically-relevant state in cells would be highly desirable.

Previous studies in other RNA viruses have identified numerous host RNA binding proteins (RBPs) that function to regulate the viral infection cycle (Li and Nagy, 2011; Ooi et al., 2019), but we are unaware of such information for SARS-CoV-2. Our group recently demonstrated that RNA structural data can be assessed using cutting-edge deep learning techniques to construct neural network models that integrate *in vivo* RBP binding and RNA features in matched cells, yielding accurate predictions about *in vivo* RBP-virus genome binding interactions (Sun et al., 2020). Given the centrality of such interactions to understanding how viruses engage with their host cells, a large-scale survey and/or prediction to determine which host RBPs interact with SARS-CoV-2 genomic RNAs during infection would be an extremely rich resource for molecular insights.

In the present study, we first investigated the *in vivo* and *in vitro* RNA secondary structures of SARS-CoV-2, as well as the structures of the untranslated regions (UTRs) structures of six related coronaviruses using a high-throughput technology known as *in vivo* click selective 2-hydroxyl acylation and profiling experiment (icSHAPE) (Spitale et al., 2015). We experimentally determined structural maps of the SARS-CoV-2 genome in infected human cells, and also characterized the *in vitro* refolded RNA structures of SARS-CoV-2 and 6 other coronaviruses. Based on the *in vivo* structural data, we then used our deep learning tool to accurately predict several dozen functionally related host cell proteins that bind to the SARS-CoV-2 RNA genome (Sun et al., 2020); among these host proteins many are vulnerable drug target.

## RESULTS

### icSHAPE-based determination of the SARS-CoV-2 RNA genome structural landscape

To delineate the genome-wide structure of SARS-CoV-2 in infected cells, we performed *in vivo* click selective 2-hydroxyl acylation and profiling experiments (icSHAPE) to assess structural information for every nucleotide of the RNA genome (Figure 1. STAR Methods). Specifically, after infecting Huh7 cells with SARS-CoV-2 for 30 hours, we treated cells with the icSHAPE reagent NAI-N3, which permeates cells, selectively reacts with single stranded nucleotides, and introduces a modification to the 2’-OH group of the sugar ring. We then purified the total RNA and performed reverse transcription, followed by cDNA sequencing library construction. Note that NAI-N3-modifications which tag single-stranded nucleotides functionally block reverse transcriptase, thus enabling detection of single-stranded nucleotides based on deep sequencing and bioinformatics analysis. Finally, we obtain an icSHAPE reactivity score for each nucleotide of the viral RNA genome (and the host cell transcriptome); scores are between 0 and 1, with higher scores indicating that a nucleotide is more likely single-stranded. Thus, based on mapping of the single-stranded nucleotides, icSHAPE analysis enables elucidation of the structural landscape of all RNAs *in vivo.*

**Figure 1.**
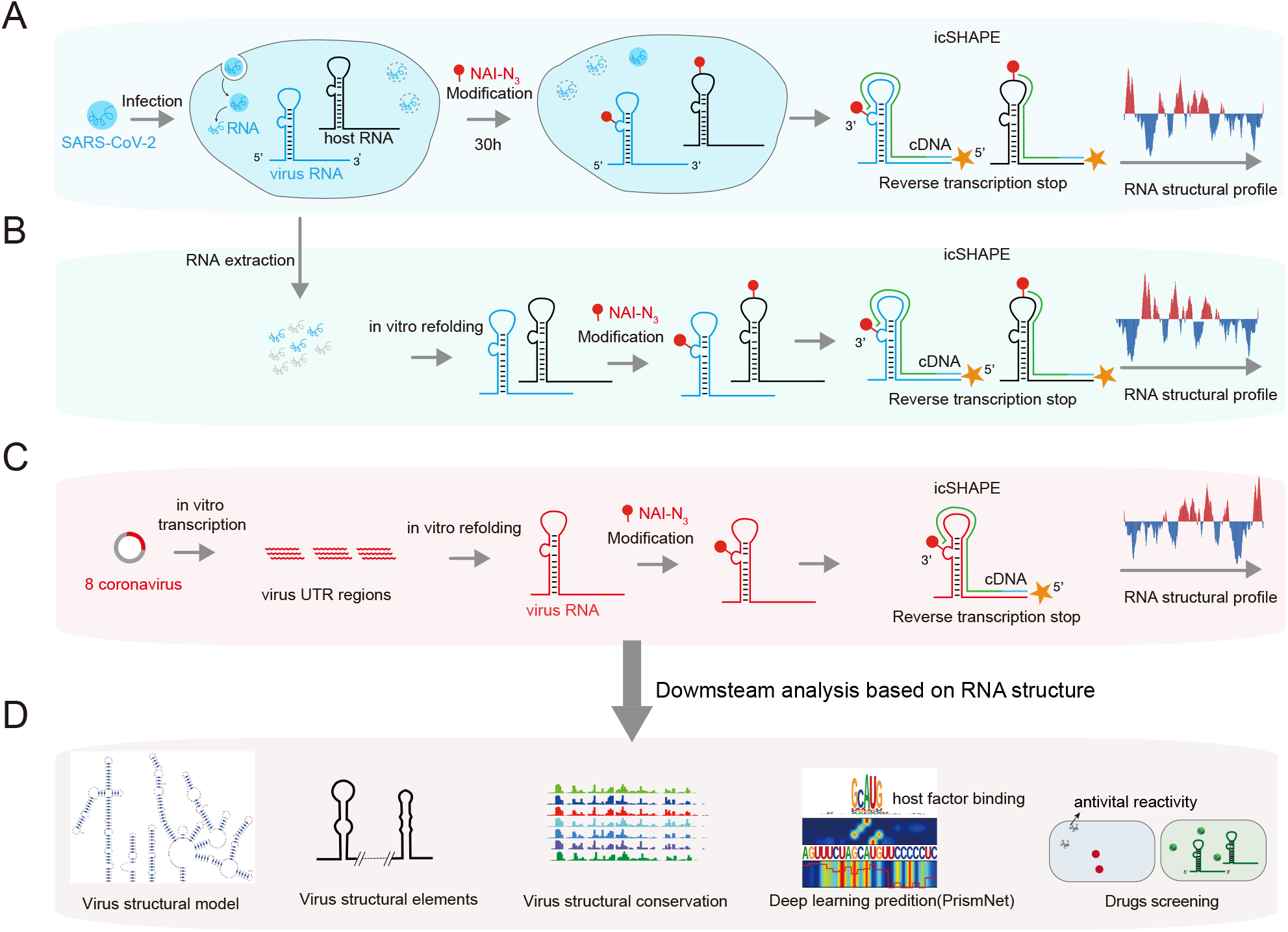
icSHAPE-based analysis of the RNA genome structure of SARS-CoV-2 and 7 other coronaviruses. **(A)** Schematic illustrating use of icSHAPE for *in vivo* studies of SARS-CoV-2 RNA genome structure. We infected Huh7 cells with SARS-CoV-2, treated these cells with the RNA structure probing reagent NAI-N3, and then performed icSHAPE experiments to characterize the *in vivo* SARS-CoV-2 RNA genome structure in host cells. **(B)** Schematic for *in vitro* structural analysis of the SARS-CoV-2 RNA genome purified from infected cells. SARS-CoV-2 RNA was purified from infected Huh7 cells, followed by *in vitro* refolding, NAI-N3 modification, and icSHAPE experiments. **(C)** Schematic for the structural characterization *in vitro* transcribed viral RNA fragments for SARS-CoV-2 RNA and 6 additional coronaviruses (*e.g.*, SARS, MERS, etc.). **(D)** Using the icSHAPE RNA structural profile data, we conducted multiple downstream analyses, including construction of *in vivo* SARS2 RNA genome structural models (based on the structural profiles as constraints), exploratory structural comparisons among coronaviruses of different subfamilies, deep-learning-based predictive analysis (using PrismNet) to identify candidate host cell proteins (host factors) impacting SARS-CoV-2 infectivity, and an infection system for drug discovery and evaluation against SARS-CoV-2.

For the *in vivo* icSHAPE structural map of the SARS-CoV-2 RNA genome, we obtained an average of about 200 million reads for each library replicate (Supplementary Table S1). Underscoring the very high quality of our sequencing data, we found that the inter-replicate Pearson correlation coefficient values are higher than 0.98 for comparison of RNA expression (RPKM) levels of the host transcriptome (Figure S2A-C); and the correlation of the RT-stop-causing NAI-N3 modifications on the viral RNA genome exceeds 0.99 (Figure S2D). Finally, we obtained icSHAPE scores for more than 99.88% of the nucleotides for *in vivo* SARS-CoV-2 RNA genome structure, by using *icSHAPE-pipe* (Li et al., 2019) (Figure 2, Supplementary Table S2).

**Figure 2.**
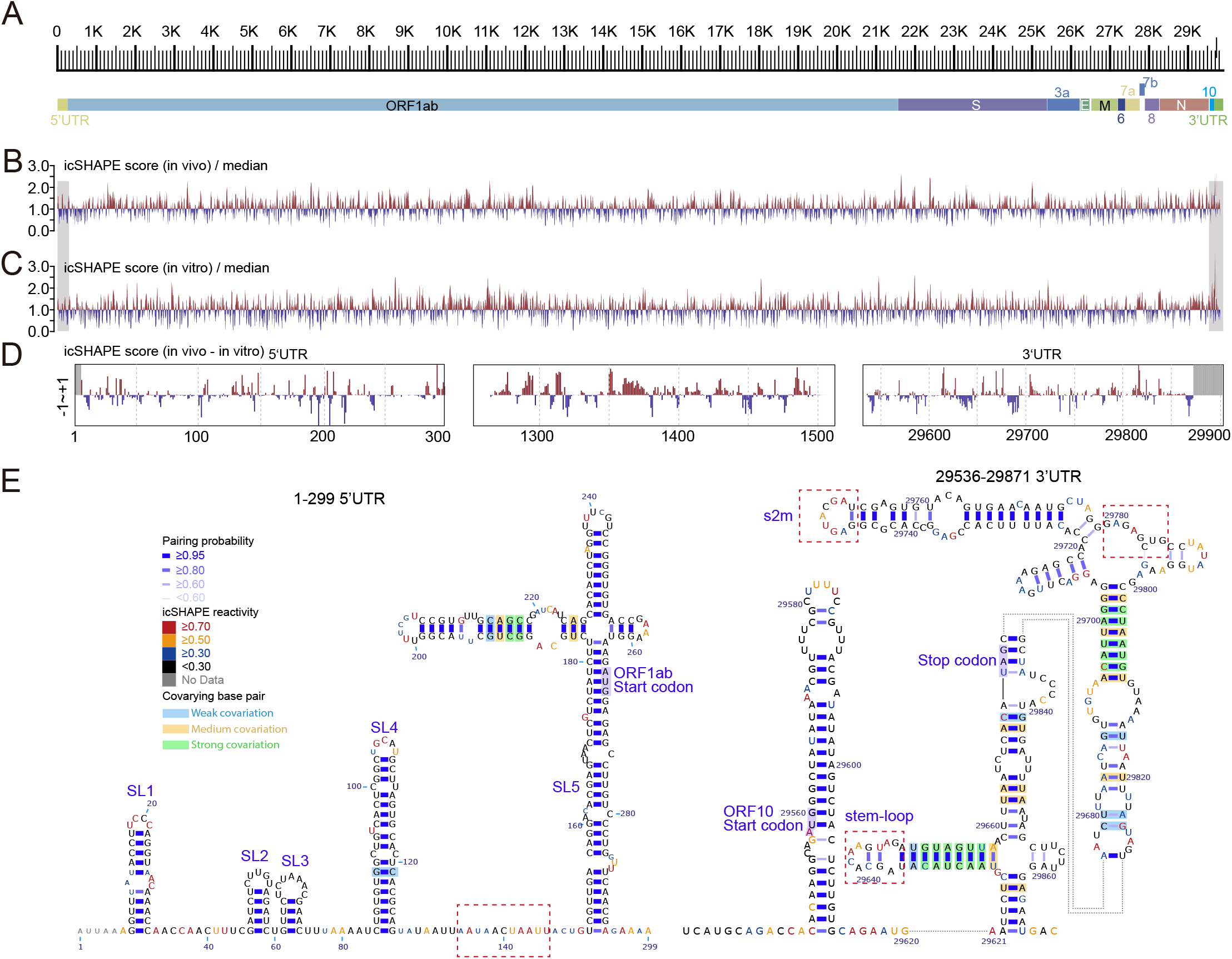
Structural Overview of the SARS-CoV-2 RNA genome. **(A)** Scale marker for the 30kb SARS-CoV-2 RNA genome (top) and a genic model showing the known organization of the genome into the 5’ UTR, the two known ORFs and nine major subgenomic RNAs, and the 3’ UTR (bottom). **(B)** Normalized icSHAPE reactivity scores across the whole SARS-CoV-2 genome based on *in vivo* data, shown relative to the global median value, with higher values corresponding to more flexible nucleotides. Blue color represents a region more likely to be in a pairing state, whereas red color represents a region more likely to be non-pairing. The normalized scores depicted here have been smoothed using a 30-nt window size. **(C)** Normalized genome-wide icSHAPE reactivity scores for the SARS-CoV-2 genome based on the *in vitro* refolding data. **(D)** icSHAPE profile of 5’UTR, ORF1ab region and 3’UTR between in *vivo* data and *in vitro* refolding data (*in vivo* – *in vitro*). **(E)** Selected RNA structural models from the 5’-UTR (left) and the 3’-UTR (right) (both with flanking regions) of SARS-CoV-2, constructed with the *RNAstructure* program using the icSHAPE reactivity scores as constraints. Nucleotides are colored by icSHAPE reactivity scores, with red and yellow colors indicating reactive nucleotides. Blue bars show base-pairs with high pairing probability. Nucleotides with a color background were predicted as covariation base-pairs.

We initially assessed the accuracy of our *in vivo* structure based on comparisons with the very recently published theoretical models of the 5’UTR and 3’UTR secondary RNA structures built from homology based on literature-annotated structures in related betacoronaviruses (Rangan et al., 2020), as well as in the Rfam database based on co-variation and pairing energy values (Kalvari et al., 2018). We built the 5’UTR and 3’UTR structure models with RNA structure modeling software tools (here we used *RNAstructure*) (Reuter and Mathews, 2010), using icSHAPE scores as constraints (Figure 2E); such methods have been extensively used and validated by other groups and ours in RNA structural studies including viral RNA structures (Li et al., 2018; Pirakitikulr et al., 2016; Watts et al., 2009) (STAR Methods). Attesting to the quality of the map generated from our *in vivo* data, the single-base resolution SARS-CoV-2 models are in strong agreement with reported structural information from predictive and *in vitro* studies (compare Figure 2E and Figure S2A-B). That is, we can easily identify the anticipated Stem-loop 1 (SL1), Stem-loop 2 (SL2), Stem-loop 3 (SL3), Stem-loop 4 (SL4), and the hyper-variable region (HVR) comprising a conserved octonucleotide sequence and the stem-loop II-like motif (s2m) in the 3’UTR (Figure 2E).

Small differences exist between our model and the Das theoretical model (shown in red dashed-line boxes, Figure 2E and Figure S2A-B), including for example the absence of a small stem-loop downstream of SL4 in the 5’UTR of our model. Our structural reactivity data based on data from Huh7 cells suggest that this region is highly single-stranded. Another difference is that we detected a stem loop structure rather than a pseudoknot in the 3’UTR of the SARS-CoV-2 genome; this alternative structural interpretation is supported by the high icSHAPE score constraint that would occur in a pseudoknot configuration. These findings highlight that *in vivo* structural information is critical for building of physiologically relevant structural models.

We also used icSHAPE to conduct *in vitro* structural analysis of the SARS-CoV-2 RNA genome. SARS-CoV-2 RNA was purified from infected Huh7 cells, refolded *in vitro*, then modified with NAI-N3, with the remaining steps and data analysis the same as *in vivo* icSHAPE (Figure 1B, Figure 2C). Similar to previous studies (Spitale et al., 2015; Sun et al., 2019), comparative analysis between *in vitro* and *in vivo* data focused on the revealed many common stable structural regions, but also indicated substantial differences, with a 0.58 Pearson correlation coefficient between the *in vitro* and *in vivo* structural profiles at the whole-genome level (Figure 2D, Figure S1E). Additionally, and consistent with findings from previous studies of viral RNA genome structure, comparisons of our models indicated that the SARS-CoV-2 RNA genome *in vivo* appears to be more single-stranded than *in vitro* (Figure S1F). These detected differences between our *in vitro* and *in vivo* structural profiles strongly emphasize the utility of studying RNA structures in their cellular context to uncover biologically relevant conformations.

### In vivo structural model of the whole SARS-CoV-2 RNA genome

We extended the validated approach of *in vivo* RNA structure modeling with icSHAPE scores as constraints to the whole SARS-CoV-2 RNA genome (STAR Methods). We tested different parameters for modeling of the SARS-CoV-2 5’UTR and 3’UTR structures, and then used the parameters which generated the most consistent structure with the Das and Rfam models to conduct genome-wide modeling. Given that RNA structural modeling is typically most successful with small RNA fragments (Li et al., 2018), we used a sliding window (window = 5000nt, step = 1000nt) strategy to more accurately model RNA structures; note that we followed a common strategy of preferencing structure models with higher pairing probabilities when modeling RNA structures of overlapping regions.

We assessed co-evolutionary evidence to support our final models based on the R-scape tool and on a previously reported pipeline to call co-variant pairs from 258 coronavirus genomes (Figure S2C. STAR Methods) (Li et al., 2018; Pirakitikulr et al., 2016). In total, we found 143 co-variant pairs, including 6 in the 5’UTR, and 12 in the 3’UTR (Figure 3, S3). The flanking regions of the UTR also contains many co-variant pairs (1 in 5’UTR flanking and 19 in 3’UTR flanking), indicating potential regulatory impacts from these detected RNA structures. Interestingly, we observed a strong combination of 5 co-variations within a duplex formed by the interaction between 3’UTR and “ORF10”, a flanking cryptic ORF. ORF10 was predicted computationally, but has not been supported with any protein-level evidence (Kim et al., 2020), and a recent study using nanopore direct RNA sequencing did not identify the predicted subgenomic RNA ORF10. Our structural data raised the possibility that this region may be a part of the 3’UTR, potentially functioning as a stable RNA structure (Figure 3).

**Figure 3.**
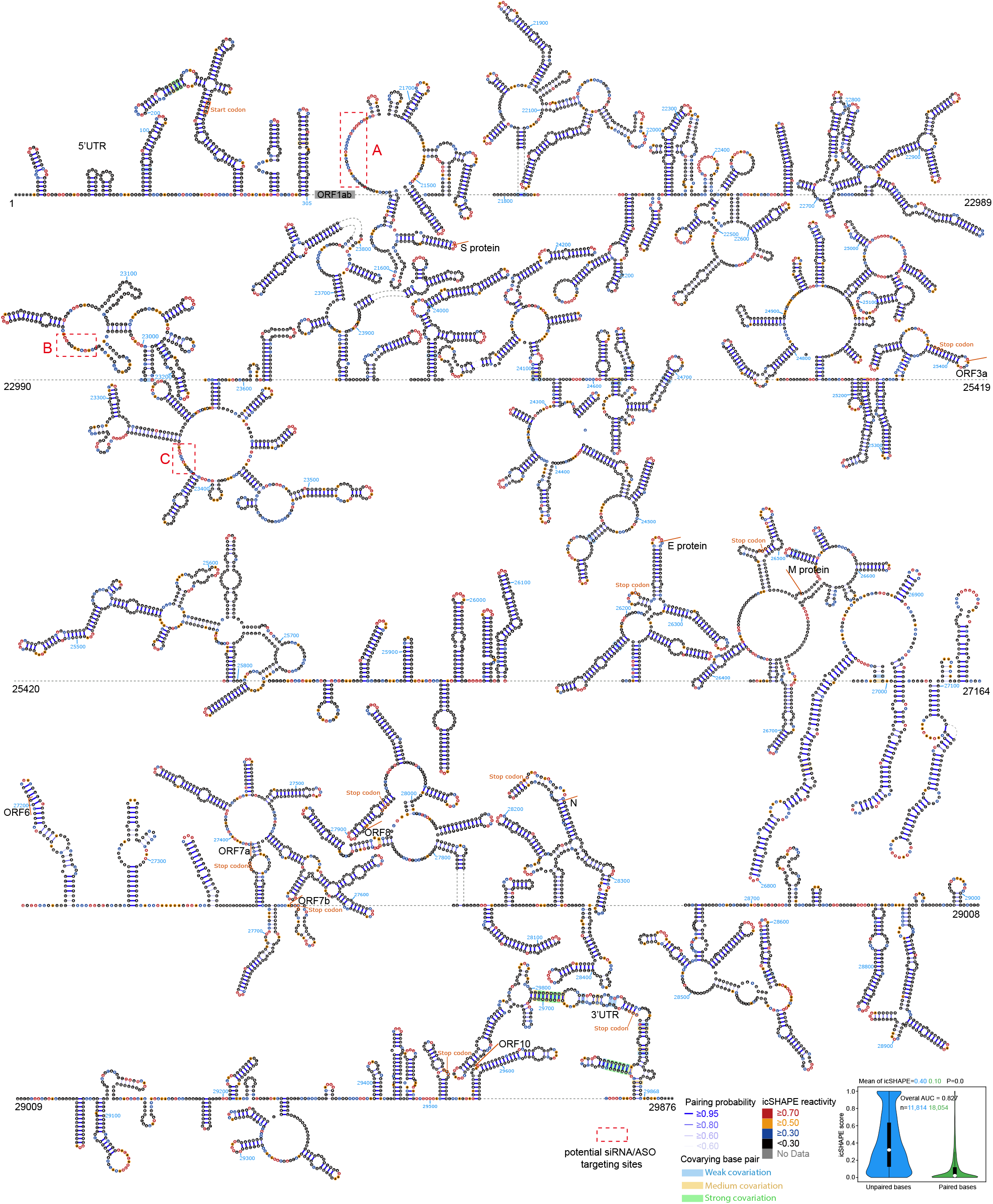
Structural model (21000nt–29903nt) of the SARS-CoV-2 RNA genome. Nucleotides are colored with icSHAPE reactivity scores; blue bars show base-pairs with high pairing probability. Nucleotides with a green background were predicted as covariation base-pairs. The boxplot insets at the bottom show the distributions of icSHAPE reactivity scores. Note that a full-length structural model of the SARS2 RNA genome is shown in the Supplementary Figure.

Our model revealed a rich abundance of long single-stranded nucleotide regions; beyond their overall structural impacts, such regions are understood as attractive targets for interventions including siRNA and antisense oligonucleotide (ASO) therapeutics. We identified 469 such regions, covering 4,863 bases from all ORFs and subgenomic RNAs of SARS-CoV-2 (Supplementary Table S3, STAR Methods). Thus, our SARS-CoV-2 RNA genome model built using icSHAPE constraints provides reliable structural information and can be understood as a valuable resource to advance our understanding of this virus and its potential vulnerabilities.

### Structural conservation and divergence across the non-coding regions of the coronavirinae family

In principle, conserved RNA structures across evolution suggest potential functional importance. To examine structural conservation of the non-coding regions in *Coronavirinae*, we performed icSHAPE-based analysis on the 5’UTR and 3’UTR (about 1000 nt in total. Supplementary S4) across 7 different coronavirus genera (Alphacoronavirus, Betacoronavirus) and lineages (lineage A, B, C D) (Figure 4A). Precisely, we examined the reference SARS-CoV-2 strain and a mutant SARS-CoV-2-241T with a C-to-T mutation at 241nt, SARS-CoV (lineage B, Betacoronavirus), MERS (MERS-CoV), NL63 (HCoV-NL63, Alphacoronavirus), HKU1 (HCoV-HKU1, lineage A, Betacoronavirus); and two bat coronaviruses: HKU5 (BtCoV-HKU5, lineage C, Betacoronavirus) and HKU9 (BtCoV-HKU9, lineage D, Betacoronavirus). Note that the 5’UTR sequences all included about 300 nt of flanking sequence. We synthesized the corresponding DNA sequences and used *in vitro* transcription to obtain the RNA fragments. Following re-folding, we applied NAI-N_3_ and then characterized their RNA structures using icSHAPE.

**Figure 4.**
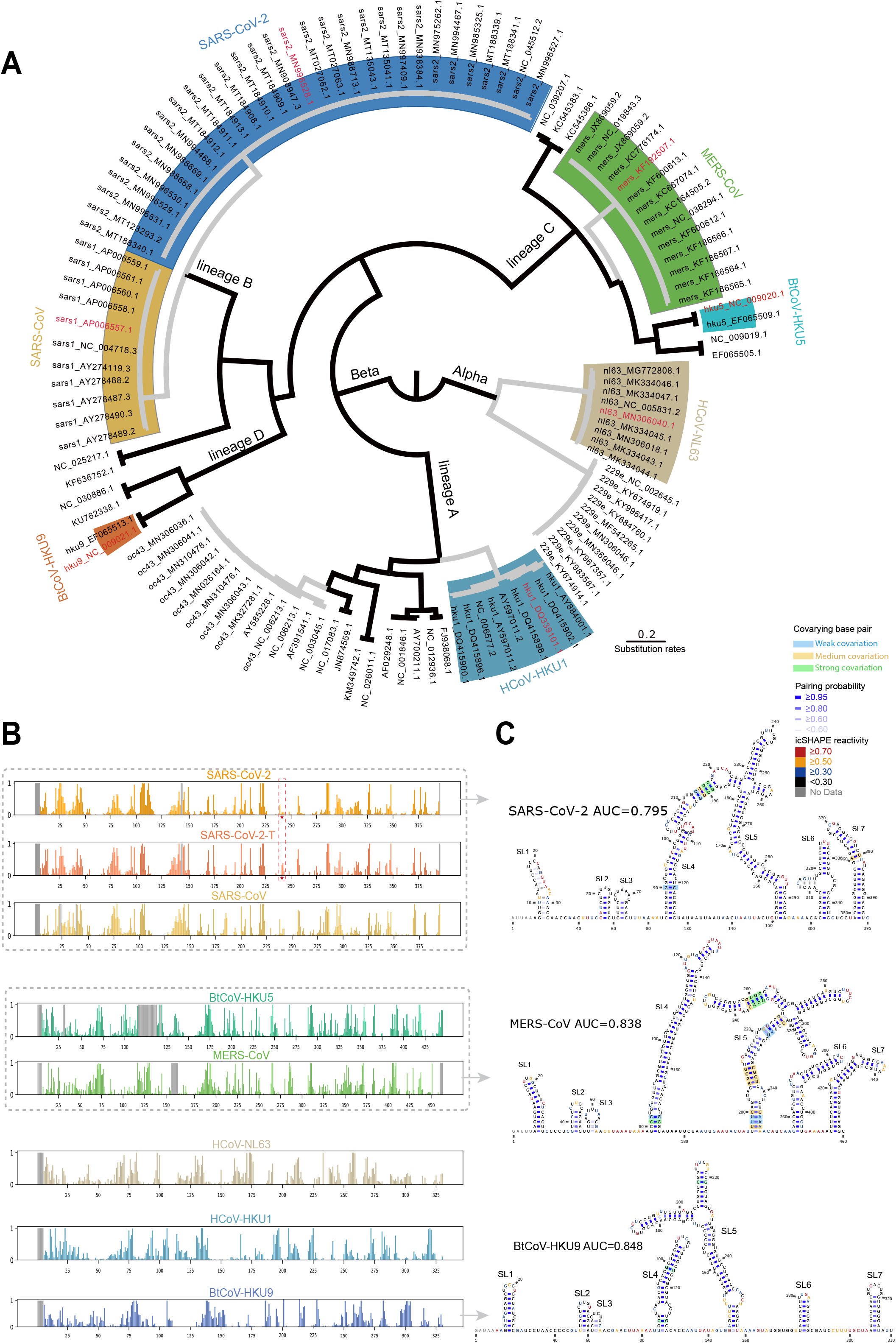
Comparative analysis reveals structural characteristics and conservations among 5’-UTRs of 7 coronaviruses. **(A)** Phylogenetic diagram showing the evolution of the coronavirinae Alpha and Beta subfamilies. The 7 coronaviruses are from the Alpha coronavirus subfamily (HCoV-NL63) and different lineages of the Beta coronavirus subfamily, including lineage A (HCoV-HKU1), lineage B (SARS-CoV, SARS-CoV-2), lineage C (MERS-CoV, BtCoV-HKU5), and lineage D (BtCoV-HKU9). Virus strains investigated with icSHAPE analysis are colored in red. **(B)** icSHAPE reactivity scores for the 5’-UTRs (with flanking regions) for the selected coronaviruses. Viruses with sequence identify higher than 70% are clustered into the same group. **(C)** Structural models of the 5’-UTRs of SARS-CoV, SARS-CoV-2 and BtCoV-HKU9. Nucleotides are colored with icSHAPE reactivity scores, blue bars show base-pairs with high pairing probability, and nucleotides with color backgrounds were predicted as covariation base-pairs.

Of particular note, a recent study found that a SARS-CoV-2 variant with a glycine at the residue 614 of the Spike protein may confer a fitness advantage; this variant has become the dominant pandemic form of the virus (Korber et al., 2020). The study’s authors suggested that this mutation may facilitate cell entry or reduce immune response to increase overall transmission rates. Interestingly, this mutation is almost invariably accompanied by a 241 C-to-T mutation in the SARS-CoV-2 5’UTR. We therefore included the SARS-CoV-2-241T mutant in our comparative structural analyses, and we did observe a modest structural change around this position (Figure 4B), suggesting that this non-coding mutation may also contribute to an increase in transmission.

Over all the sequences we examined, icSHAPE profile data for these diverse noncoding genome regions revealed conserved structures characteristic of the various genera and linages, largely consistent with the phylogeny (Figure 4B), both in 5’UTR (Figure S4) and 3’UTR (Figure S5). For example, the 5’UTRs for all of the lineage B group members (SARS-CoV and SARS-CoV-2) each contain 7 stem-loop RNA structures, and these structures occur in the same order (Figure 4C, S4A). Remarkably, lineage C group (MERS-CoV and BtCoV-HKU5) also contain seven almost identical stem-loops, again in the same order, even though these viruses have a sequence similarity of only 46.5%~47.3% with lineage B (Figure S4B). The more distant lineage D betacoronavirus (BtCoV-HKU9) also contains seven similar stem-loops, despite having only 39.3~40.7% sequence similarity with lineage B and C (Figure S4B). Nevertheless, lineage D virus has slightly longer SL6 and SL7 than lineage B and C (Figure 4C, S4A). Notably, although some theoretical models based on co-variation show similar structural architecture (Figure S4C), others are very different and miss conserved structural elements (Figure S4D, MERS-CoV, BtCoV-HKU9).

Strikingly, despite having similar levels of sequence similarity to betacoronaviruses of lineages B, C, and D, the lineage A betacoronavirus (HCoV-HKU1) and Alphacoronavirus (HCoV-NL63) (37.5%~ 46.3% in 5’UTR and 35.0%~45.2% in 3’UTR) each form very different structures, although all share stem-loop SL1(Figure S4B, S4E and S5B). This structural divergence suggests that the non-coding regions of these viruses may exert distinct functions and may be subject to different forms of regulation. Our characterization of structural features from 7 coronavirus lineages in combination with our phylogenetic analysis revealed how coronavirus evolution has resulted conserved as well as divergent RNA structures.

### In vivo RNA structure predicts translation efficiency and species abundance of subgenomic RNA

RNA structure affects the localization, splicing, translation efficiency (TE), and stability of cellular RNA on a genome-wide scale (Bevilacqua et al., 2016; Mortimer et al., 2014; Piao et al., 2017; Wan et al., 2011). Previous studies have reported positive correlations between translation efficiency and the frequency of single-stranded regions in the 5’-UTR of a given RNA transcript (Beaudoin et al., 2018; Mustoe et al., 2018). SARS-CoV-2 generates a number of subgenomic viral RNA sequences which encode important structural proteins including the spike protein, the envelop protein, the membrane protein, the nucleocapsid protein, and other accessory proteins (Figure S6A) (Kim et al., 2020). By correlating our *in vivo* SARS-CoV-2 RNA structure data with translation efficiency (TE) data of subgenomic viral RNAs reported in a recent study (Bojkova et al., 2020), we found a high Pearson correlation coefficient between TE and RNA structural features (r = 0.717, p =0.029, Figure S6B). That is, for SARS-CoV-2 RNAs in cells, the TE is generally higher for RNA molecules of more single-stranded 5’-UTR5’-UTR.

We also found that the abundance of a given subgenomic viral RNA is positively correlated with the frequency of single-stranded regions present in its structure. For context, subgenomic viral RNAs are generated from the minus-strand of viral RNA intermediates that are synthesized via a process known as “discontinuous transcription”. In this process, transcription begins at the 3’ end of the viral RNA genome and halts upon reaching a so-called TRS-B sequence—a transcription-regulatory sequence in the main body of the molecule positioned adjacent to the ORFs; this is followed by resumption of transcription upon switching the template to the 5’ TRS-L (TRS in the leader) sequence. The intact minus-strand RNA intermediates form from the fusion of TRS-B and TRS-L (along with the sequence uptream of TRS-B), and then serve as the templates of positive-sense subgenomic viral RNAs generation (Figure S6A).

A recent study reported quantification of the SARS-CoV-2 subgenomic viral RNA population based on long-read sequencing (Kim et al., 2020). Using this abundance data alongside our structural data, we detected a positive correlation between the abundance of a particular subgenomic viral RNA and the level of single-stranded regions positioned within its 5’ TRS-B region (Figure 5A, left, r = 0.24, p = 0.035). Pursuing this, we re-examined our sequencing data from the initial icSHAPE analysis to accurately calculate RNA structure scores for the TRS-L region of each subgenomic viral RNA, specifically by identifying and exclusively counting those reads that i) cross a fusion site and ii) specifically map to a confirmed subgenomic viral RNA (Figure S6A, left, STAR method). Again, subgenomic viral RNAs if more single-stranded TRS-L were much more abundant (Figure 5A, right, r = 0.43, p = 0.004. See an example of N and pp1ab in Figure 5B),supporting that the overall abundance of a particular viral RNA molecule is strongly impacted by its RNA 5’ structure, presumably because of differential impacts on discontinuous transcription.

**Figure 5.**
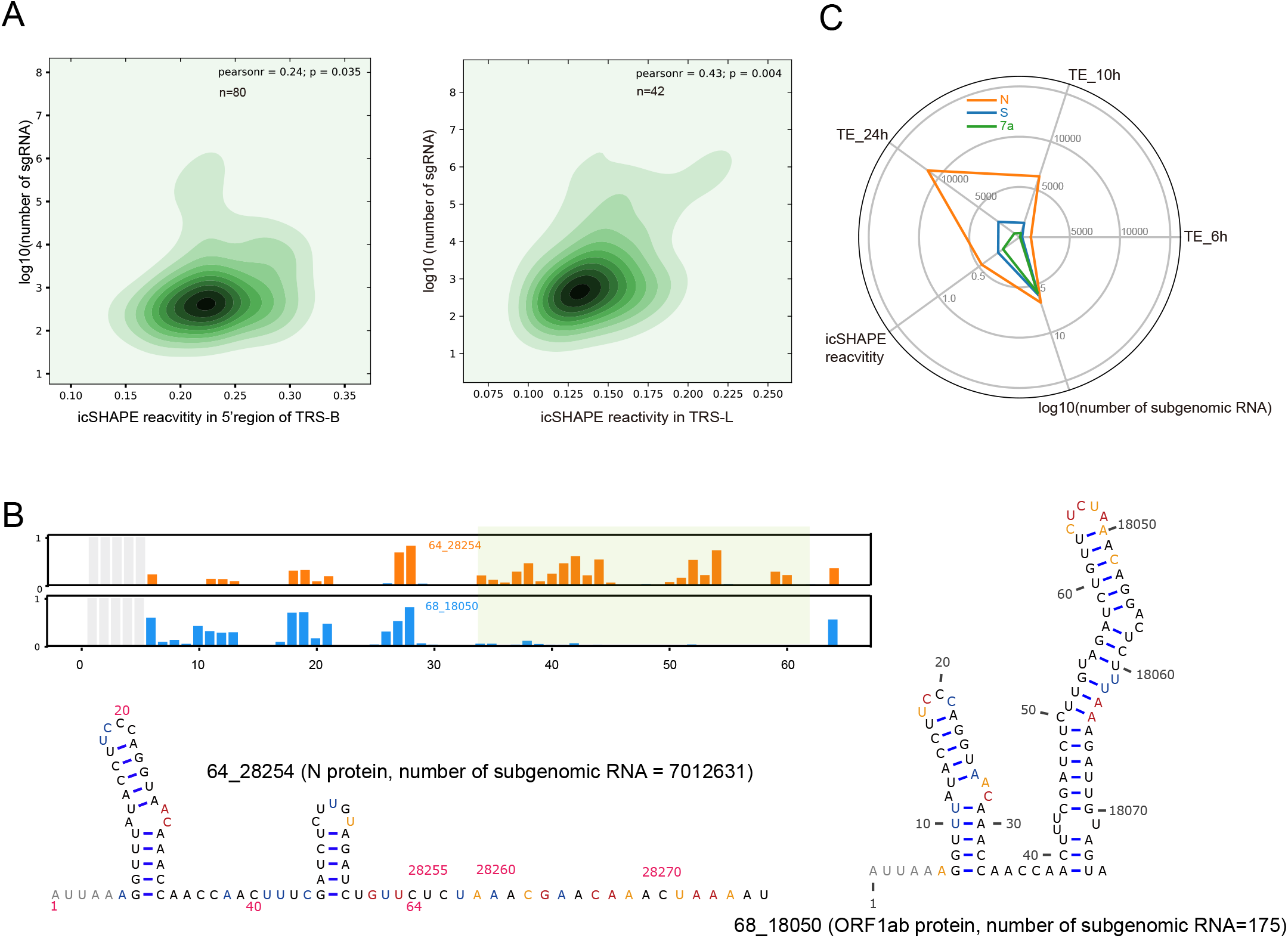
RNA structure functionally impacts both the generation and translation of SARS-CoV-2 subgenomic RNAs. **(A)** Scatter plots showing correlation values between the detected abundance of a given subgenomic RNA versus its icSHAPE reactivity score. **(left)**shows the correlation for the specific regions of the 5’-UTR of each subgenomic RNA. **(right)**shows the correlation for the common region of the 5’-UTR of all subgenomic RNAs, *i.e.*, the TRS-L region. The two-tailed P value was calculated using the Python package function scipy.stats.pearsonr. *r* is the Pearson correlation efficient. **(B)** The icSHAPE profile and structural models of the TRS-L region of the subgenomic RNAs of nucleocapsid (N) and polypeptide 1b (pp1b), predicting the relative abundance of the subgenomic RNAs. RNA structural models were here predicted using the icSHAPE score constraint, as above (see Fig 2E). **(C)** Radar diagram showing the correlation of 5′-UTR RNA structure, translation efficiency at different time points post viral infection (data from Munchet et. al. 2020), and the number of subgenomic RNAs for the nucleocapsid protein (N), the spike protein (S), and the accessory 7a protein (data from Kim et. al. 2020).

The coupling of transcription and translation of cellular RNAs is an important phenomenon in gene expression, fulfilled through different mechanism of regulation, including m6A, RNA polymerase II subunits and promoter sequences (Harel-Sharvit et al., 2010; Slobodin et al., 2017; Zid and O’Shea, 2014). In our previous study, we established RNA structure as a link in connecting transcription and translation. In our present analysis of SARS-CoV-2 subgenomic RNAs, we also observed apparent impacts of RNA structure on the coordination of subgenomic viral RNA generation and translation (Figure 5C). Collectively, our *in vivo* RNA structure predicts and connects the abundance and the translation efficiency of SARS-CoV-2 subgenomic RNAs.

### PrismNet accurately predicts host proteins that bind to the SARS-CoV-2 RNA genome based-on in vivo RNA structure using deep learning

Host cell RNA binding proteins (RBPs) play essential roles in regulating virus translation, replication, and degradation (Li and Nagy, 2011; Ooi et al., 2019). Thus, characterizing interactions between RBPs and viral RNA is a is fundamental step for understanding the infection process and for identifying potential targets to enable development of innovative antivirus therapies. Although genome-scale viral RNA genome data is relatively scarce and methods to predict and detect viral RNA genome-host RBP interactions are only in their infancy, it is notable that previous studies have reported functional interactions between RBPs and the 5’UTRs of coronavirus genomes (Sola et al., 2011b). Our group has recently developed a tool— PrismNet (<u>P</u>rotein-<u>R</u>NA Interaction by <u>S</u>tructure-informed <u>M</u>odeling using deep neural <u>Net</u>work)—which we here employed to predict host RBPs and their putative RNA genome binding sites (Sun et al., 2020). Very briefly, PrismNet constructs and trains a deep neural network to model the interactions between an RBP and its RNA target by integrating big data from *in vivo* RBP binding assays and RNA structural analyses obtained from matched cellular conditions.

We applied PrismNet to predict RBP binding on the SARS-CoV-2 RNA genome, and identified 40 and 43 host proteins that bind to the 5’UTR and 3’UTR, respectively (Figure 6A, Supplementary Table S5). Several of our predicted proteins were previously reported to directly or potentially binding with RNA of other coronaviruses. For example, the heterogeneous nuclear ribonucleoprotein hnRNPA1 was shown to bind with viral RNA and impact RNA synthesis for mouse hepatitis virus (MHV) (Shi et al., 2000). PTBP1 was reported to bind to transmissible gastroenteritis virus (TGEV) RNA and is involved in viral gene expression (Sola et al., 2011a). The NPM1 protein interacts with the nucleocapsid protein of SARS-CoV, which affects viral particle assembly (Zeng et al., 2008). Although there are no studies reporting any association of the autoimmune-disease-related protein TROVE2 (Reed and Gordon, 2016) with coronavirus infection; however, we found that TROVE2 shows high affinity (0.979) with SARS-CoV-2 RNA based on our prediction, identifying it as a candidate host RBP to help study how SARS-CoV-2 infection may deleteriously impact host immune responses.

**Figure 6.**
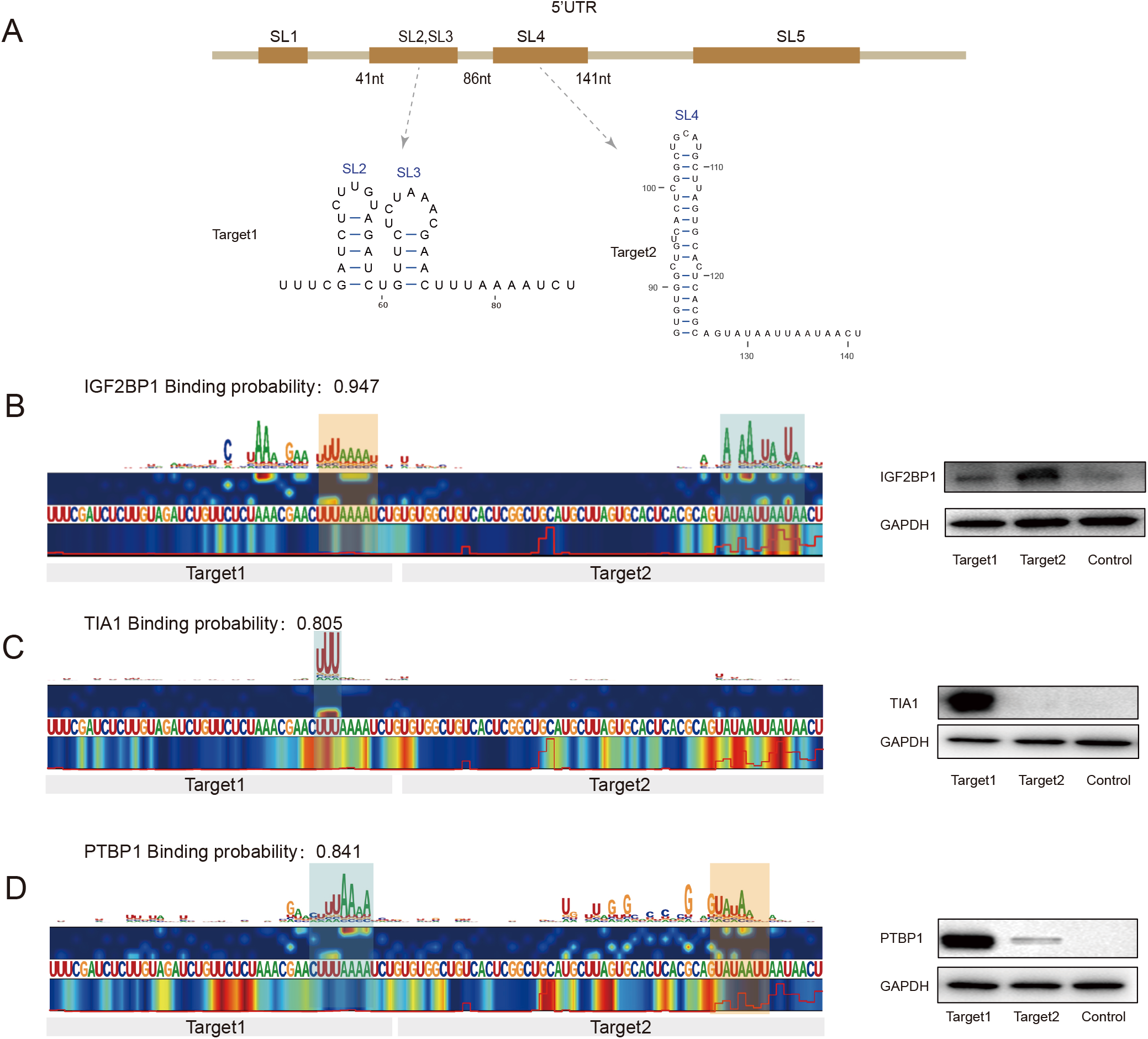
Applying deep learning to the in vivo SARS-CoV-2 RNA structure accurately predicts host proteins that bind to SARS-CoV-2. **(A)**Schematic for the 5′-UTR of SARS-CoV-2, including five stem-loop (SL) regions. Two RNA structures, including a combined stem-loop 2 (SL2) and stem-loop 3 (SL3) as well as stem-loop 4 (SL4), were synthesized for validation experiments (see below). **(B-D)**Left: Saliency maps from PrsimNet showing the predicted binding site of the RNA binding proteins IGF2BP1 (B), TIA1 (C), and PTBP1 (D). Grey bars indicate the range of synthesized RNA fragments. Green color shows predicted strong binding sites, while pink color shows sites predicted to have relatively weaker binding, *with predicted binding probabilities shown at the top*. Right: western blotting for the RNA pull-down assay based on the synthesized RNA fragments (SL2-SL3, SL4).

Inhibition of stress granules affects the replication of MERS-CoV, and our analysis predicted that several stress-granule-related proteins such as TIA1(Onomoto et al., 2014), IGF2BP1, and ELAVL1 (Markmiller et al., 2018) also showed high affinity with SARS-CoV-2 RNA. These multiple predictions collectively indicate that SARS-CoV-2 infection may somehow inhibit formation of stress granules in host cells, perhaps in a manner similar to MERS-CoV’s hijacking of related RBPs. We therefore focused on IGF2BP1, TIA1, and PTBP1 as a potential host factors for SARS-CoV-2 infection to validate our prediction data. We synthesized 2 host sequences which include the SL2, SL3 fragment (Target 1) and the SL4 fragment (Target 2) to use in RNA pull-down assays (Figure 6A). PrsimNet predicted IGF2BP1, TIA1, and PTBP1 bound to SARS-CoV-2 RNA (with binding probabilities: 0.947, 0.805 and 0.841). Our pull-down assays showed that the detected binding strengths correlated well with the predicted binding scores based on the saliency map from PrismNet (Figure 6B-D, left). Specifically, TIA1 only bound to Target1, whereas PTBP1 also exhibited very weak affinity for Target2. IGF2BP1 can bind to both of the Target1 and Target2, albeit with stronger affinity for Target 2 (Figure 6B-D, right). Our empirical data thus support that PrimNet can accurately predict host proteins that bind to the SARA-CoV-2 RNA genome, opening the door for multiple hypothesis-driven studies to determine whether and how such binding interactions with these host RBPs may regulate viral infectivity.

## DISCUSSION

In this study we experimentally determined structural maps of the SARS-CoV-2 genome in infected human cells, as well as the structure for *in vitro* refolded RNA of SARS-CoV-2 and 6 other coronaviruses. Our host cell data confirms the presence of stable, conserved structural elements from theoretical analysis, while also revealing many previously unknown structural features that apparently affect viral translation and discontinuous transcription in cells. Based on our *in vivo* structure data, we then accurately predicted several dozen functionally related host cell proteins that bind to the SARS-CoV-2 RNA genome by using a deep learning neural network, and finally showed that some of these host proteins are vulnerable drug targets for reducing SARS-CoV-2 infection.

In addition to encoding viral proteins, the SARS-CoV-2 RNA genome itself functions as a molecular hub to interact with many cellular factors, presenting multiple levels of complexity for the regulation of viral infection and disease. As discovered previously for many other viruses, including HIV (Watts et al., 2009), HCV (Pirakitikulr et al., 2016), dengue (Dethoff et al., 2018), and ZIKV (Li et al., 2018), much of the regulation and function of RNA viral genomes is mediated by higher-order RNA structures. For coronaviruses, studies have also identified different RNA structure elements that functionally impact viral life cycles. Most coronavirus 5’ UTR’s harbor a number of loops, with many showing heightened sequence conservation across betacoronaviruses, and various stems demonstrating functional roles in viral replication. For example, studies suggested that the first stem-loop (SL1) the in 5’UTR is necessary for viral replication (Li et al., 2008). Flanking the 5’UTR of SARS-CoV-2, the third stem-loop contains a TRS core sequence (CS region, CUAAAC), and PrismNet predicted that some RBPs bind with this genome structure (Figure 6), suggesting that it is critical for discontinuous transcription characteristic to coronaviruses. In viral genome 3’UTRs, mutually exclusive RNA structures have been shown to control various stages of the RNA synthesis pathway. Recent virus structural modeling efforts using SARS-CoV-2 genome sequences have confirmed the existence of many of these stem-loops and driven predictions of yet more of these in SARS-CoV-2 (Andrews et al., 2020; Rangan et al., 2020).

Our work emphasized that most stem-loops exist in both re-folded RNA molecules and in viruses within host cells, implying that these robustly auto-generating structures are very likely to exert functional impacts. Interestingly, we observed that the proposed loop region in SL3 is not reactive, supporting the possibility of long-range functional interactions with downstream TRS-B regions, which is understood as integral for successfully discontinuous transcription (Enjuanes et al., 2006). We also noticed that the small stem-loop downstream of SL4 proposed by the Das model is absent from our *in vivo* structural data; rather than a stem-loop, our results indicate this regions adopts a long, single-stranded conformation *in vivo*; interestingly, the sequence context of this region is AU-rich, suggesting it may be a hotspot for the binding of RBPs that prefer AU-rich single-stranded structure elements.

Guided by recent epidemiological reports, we analyzed the C-to-T mutation at the 241 nt, which is positioned in the loop region of SL5. This mutation both creates a poly-U sequence motif and modestly increases its preference to adopt a single-stranded conformation, thereby potentially creating an attractive binding site for RBPs that bind to single-stranded poly-U sequences. Our findings thus offer structural insights to help better understand how this now-prevalent SARS-CoV-2-241T allele is contributing to increased rates of transmission.

Beyond confirming the presence of a long, single-stranded region between SL3 and SL4, our study identified many more such regions in the SARS-CoV-2 genome. These regions should be understood as potentially attractive target sequences for interventions using technologies like siRNA, ASO, and sgRNA. Importantly, our work also revealed structural elements having strong co-evolution support, which usually suggests stable, functionally conserved RNA structures, throughout the genome (including in CDS regions). Computational methods like ROSETTA and FARFAR have become efficient for tertiary structure modeling when accurate secondary structural models are available. It is thus possible to build reliable tertiary structure models for some parts of the SARS-CoV-2 genome; such efforts may reveal druggable pockets vulnerable to small molecules. Indeed, there are previously reported examples for specifically targeting such functional RNA structural elements to disrupt viral infectivity (Ren and Patel, 2014), so the elucidation of RNA structures can facilitate target discovery and the development of antiviral therapeutics.

Our *in vivo* RNA structure provides a groundwork to accurately predict host RBPs that bind to SARS-CoV-2 genome, as we have demonstrated recently in different cellular contexts. We used a deep learning method, PrismNet, trained on more than 1594 binding sites together with *in vivo* RNA structures obtained from matched cell lines for 99 RBPs. Our application of PrismNet here strongly supports the accuracy of its predicted interactions. In addition to recruiting the necessary translation machineries, SARS-CoV-2 potentially interacts with many RNA metabolism proteins including helicases based on our prediction (such as DDX42), which potentially are hijacked by the virus to help evade cell innate immune response (Beachboard and Horner, 2016). In particular, the results also revealed a few stress granule proteins interacting with SARS-CoV-2 genome (Such as TIA1). Previous study has identified West Nile virus minus-strand 3’ terminal stem loop (SL) RNA specifically interact with stress granule compartments TIA1 and facilitates flavivirus genome RNA synthesis, also inhibits SG formation(Emara and Brinton, 2007). The information of how SARS-CoV-2 RNA genome interact with host cellular proteins has been extremely limited by now. Our results thus provide a rich resource to understanding the molecular mechanism of viral infection.

The identification of host RBPs that bind to RNA genomes can be exploited as a unique opportunity for the effective development of antiviral drugs. For example, drugs targeting DDX42 may inhibit the function of DDX42 that SARS-CoV-2 are more sensitive to innate immune system. Certainly, many more studies will be required before any of the drugs identified by our study can be deployed as therapeutics for treating COVID-19. But overall, the data in our study clearly has great promise for the repurposing of existing drugs and the identification of new agents that target the *in vivo* viral RNA structure and host RBPs to fight against the still-ongoing pandemic and to combat viral disease more generally.

## Acknowledgments

We thank members of the Zhang lab for discussion. We thank Professor Jianbin Wang for the help of the project. This work is supported by the National Natural Science Foundation of China (Grants No. 31671355, 91740204, and 31761163007), National Key R&D Program of China (2020YFA0707600), and Tsinghua-Cambridge Joint Research Initiative Fund. We thank the Tsinghua University Branch of China National Center for Protein Sciences (Beijing) for the computational facility support. L.S. was supported by the Tsinghua-Peking Center for Life Sciences Postdoctoral Fellowship.

## Author Contributions

Q.C.Z. conceived the project. Q.C.Z. and J.W. supervised the project. J.R. and L.R. prepared the SARS-CoV-2 virus. L.S. performed icSHAPE experiments and validation experiments. K.X. and W.H predicted the RBP binding by the deep learning models. P.L., L.S. and X.Z. analyzed all the results. X. J., T.X., S.Z., and Q. D assisted with analysis. Q.C.Z.and L.S. wrote the manuscript with inputs from all authors.

## Declaration of Interests

The authors declare no competing interests.

## Notes

### Competing Interest Statement

The authors have declared no competing interest.

